# Ecological coherence in abundance dynamics across terrestrial and marine assemblages

**DOI:** 10.1101/2025.11.05.686795

**Authors:** Alexandre Fuster-Calvo, Katherine Hébert, Aleksandra J. Dolezal, Christine Parent, François Massol, Mia Tayler Waters, Dominique Gravel

## Abstract

Understanding how communities respond to environmental change requires assessing not just overall variability but also the structure of co-variation among taxa. We frame this idea under the Ecological Coherence (EC) framework, which generalizes previous notions such as community synchrony or coherence. EC captures the structure of co-responses among taxa within assemblages and can be expressed through two complementary objects: (1) the co-response matrix *C*, which contains all pairwise correlations between taxa and can be used to identify clusters of taxa with coherent responses as well as the contributions of individual species to community-wide coherence; and (2) the *EC distribution*, which summarizes the overall profile of co-responses by capturing their shape, spread, and central tendency across the community. By combining these two views, EC moves beyond single summary metrics and provides a richer picture of how coherence is organized within communities. Analyzing the EC distribution across 341 terrestrial and 105 marine assemblages worldwide, we found a general prevalence of weak correlations and a few strong, directional correlations. We also found that it varies with community composition, sampling effort, and biogeographic region. Moreover, the *C* matrix consistently identified a small subset of taxa with strong correlations to many others, suggesting a promising path to detecting those that may play central roles in amplifying or buffering community responses to environmental change. Our findings on EC pave the way for deeper investigations into what drives the diversity of ecological responses to environmental changes and how it shapes community dynamics, while also underscoring the need for strategically distributing ecological monitoring across trophic guilds and functional roles.

## Introduction

One of the most pressing challenges in biodiversity monitoring is upscaling the diversity of taxon-level responses to global change, such as shifts in abundance, distribution, and phenology, to the community level response (Magurran et al. 2010, Van der Putten et al. 2010, Devictor et al. 2012, Dansereau et al. 2025). While taxa experience environmental pressures individually, they do so within a network, where multiple biotic interactions shape and constrain their responses (Cosmo et al. 2023). These interdependencies mean that capturing and predicting communities under environmental change requires moving beyond taxon-level assessments and considering emergent properties at higher levels of organization (Berg et al. 2010, Trisos et al. 2020).

A key aspect of these emergent properties is how taxa within a community respond to shared environmental fluctuations. A well-known mechanism driving co-variations is the Moran effect (Moran 1953, Hansen et al. 2020), whereby distant populations exhibit correlated responses to common environmental variations. Depending on the magnitude and direction of these responses, communities can display more or less similar co-responses, which in turn influence ecosystem functioning (Kent et al. 2007). Biotic interactions may further shape how taxa co-respond within a community. Predator-prey dynamics, mutualistic dependencies, and competition introduce additional layers of complexity, leading to indirect effects that can amplify or dampen the impact of environmental change (Seybold et al. 2022). The extent to which taxa exhibit similar or divergent responses is often captured by the covariance structure of their abundance time series (Houlahan et al. 2007), spatial range (Carroll et al. 2024), or traits (Kharouba et al. 2018).

Co-responses have been investigated at different time scales and with various methodologies. Covariances in raw abundance trajectories reflect similarities in overall population trends (Larsen et al. 2025); correlations of latent variables from hierarchical models capture shared underlying processes (Warton et al. 2015); correlations of growth rates emphasize whether taxa fluctuate together from one time step to the next, reflecting common environmental forcing or biotic interactions (Robinson et al. 2013, Steiner et al. 2013). Other methodologies for quantifying co-responses include variance-ratio approaches that summarize synchrony at the community level (Loreau & de Mazancourt 2008), and phase- or frequency-based methods that capture coherence in population cycles (Cazelles & Stone 2003). Studies often refer to these approaches collectively as synchrony (Gouhier et al. 2014) and are usually performed for pairs of taxa. Here, we adopt the broader term Ecological Coherence (EC) to denote the covariance structure of taxa responses within a community, irrespective of the particular metric, response representation or time scale.

Both environmental filtering and biotic interactions have been proposed as key drivers shaping EC (Ives et al. 1999, Hegland et al. 2009). High synchrony may reflect shared environmental dependence or functional redundancy of response traits, while asynchrony can indicate functional complementarity or buffering mechanisms (Gonzalez & Loreau 2009, Granzotti et al. 2024). EC plays a central role in describing how a community reacts to environmental changes, in both composition and structure. Research in biodiversity-ecosystem function (BEF) and population synchrony suggests that incoherence can enhance ecosystem stability by promoting compensatory dynamics (e.g., “buffering effect”: Yachi & Loreau 1999, Gonzalez & Loreau 2009, Loreau & De Mazancourt 2008, Loreau et al. 2021), whereas coherent responses can lead to destabilizing synergistic effects, increasing ecosystem vulnerability to external perturbations (Wagg et al. 2018, Granzotti et al. 2024).

However, much of what we know comes from horizontal communities (i.e., single trophic level, competitive systems), typically primary producers. In consumer-resource systems, persistence of consumers requires to track its resource; environmental change—such as warming that alters thermal tolerance, development rate, or phenology—can decouple partners and create a mismatch in which resource peaks no longer coincide with consumer demand, reducing growth and recruitment (match-mismatch hypothesis; Cushing 1990, Schweiger et al. 2008, Durant et al. 2017). When multiple trophic levels are considered, the effects of coherence become structure-dependent, varying with trophic-chain length (Wang & Brose 2018), omnivory (Polis & Strong 1996), and intraguild predation (Wang et al. 2019). Empirical studies in multi-trophic communities likewise report diverse outcomes (Srednick & Swearer 2024), including complex bottom-up cascades from plants across multiple trophic levels (Scherber et al. 2010), combined intra- and inter-guild coherence shaping fish food-web responses to reef impacts (Viviani et al. 2019), and diversity-stability relationships suggested to reflect a complex interplay of interaction types and strengths (Danet et al. 2021). This highlights gaps in our theoretical understanding of how co-responses shape complex community dynamics across ecosystems, and raises a central question: how coherent are natural communities?

Empirical assessments across broad spatial and temporal scales are essential for refining ecological theory and identifying key drivers of community coherence (Lepš et al. 2019, Granzotti et al. 2024, LaMontagne et al. 2024). Large-scale biodiversity databases that compile long-term taxa occurrence and abundance data provide an unprecedented opportunity to examine ecological coherence across ecosystems (Magurran et al. 2010, Navarro et al. 2017, Proença et al. 2017). Such analyses can reveal generalities and exceptions, offering a basis for identifying their drivers, characterizing their variability, and highlighting where critical gaps remain.

However, despite the central role of taxa covariance in understanding community co-responses (Ranta et al. 2008), most studies of community synchrony rely on summary statistics that condense this information into a single metric (e.g., Houlahan et al. 2007, Thibaut et al. 2012, Thibaut & Connolly 2013, Valencia et al. 2020, Tsang et al. 2023, Granzotti et al. 2024). While these approaches offer useful insights, they often obscure the underlying distribution of covariance frequencies, magnitudes, and directions that characterize the full structure of community coherence. A more comprehensive approach is needed to report community coherence, one that retains the rich information on the structure and distribution of community covariances while remaining interpretable across diverse ecosystems.

We present a global characterization of abundance co-responses in natural assemblages based on year-to-year changes in log-abundance (log-differences, approximating per-capita growth rates). We examine the distribution of Ecological Coherence (EC) across terrestrial and marine communities. We do so with exploration of the frequency, magnitude, and direction of taxa correlations in these growth-rate responses. This approach enables us to identify generalities and exceptions, providing a reference point for generating new hypotheses and guiding future investigations into the drivers and ecological consequences of taxa co-responses, as well as highlighting key gaps in empirical data. We hypothesize that EC distributions are structured rather than random, reflecting both evolutionary history and ecological context. Specifically, we predict that (1) EC distributions differ across biomes; (2) variation in pairwise coherence strength is partly explained by taxonomic relatedness and assemblage context; (3) some genera contribute disproportionately to EC, acting as key drivers of assemblage-level coherence; and (4) estimates of EC are sensitive to sampling completeness and temporal resolution.

## Methods

### Defining and quantifying Ecological Coherence (EC)

Co-responses quantify the extent to which taxa respond similarly or differently to environmental change—i.e., their coherence (Figure 1a)—and are typically expressed as covariances or correlations in abundance (Gouhier et al. 2014), spatial range (Carroll et al. 2024), or traits (Kharouba et al. 2018). At the community level, co-responses summarize all pairwise covariances or correlations, and most studies have reduced this information to a single index of coherence (Hallett et al. 2016). Building on this, we define Ecological Coherence (EC) as the statistical structure of abundance co-responses within a community, expressed through two related objects. The first object is the co-response matrix *C*, a symmetric matrix in which each element represents the correlation in abundance dynamics between a pair of taxa (Figure 1b). Note we refer to this as a “co-response matrix” as a general term, acknowledging that different methods could be used to compute co-responses (Gouhier & Guichard 2014). In this study, we specifically quantify co-responses in abundance dynamics using Pearson correlation coefficients between taxa’s yearly rate of change in abundance time series (see more details below). The second object, the *EC distribution*, is the distribution of values in *C*. It provides a visual profile of the direction and strength of pairwise co-responses within the community, capturing the variability of co-responses to environmental change, characterized by the shape, spread, and central tendency (Figure 1c).

**Figure 1.**
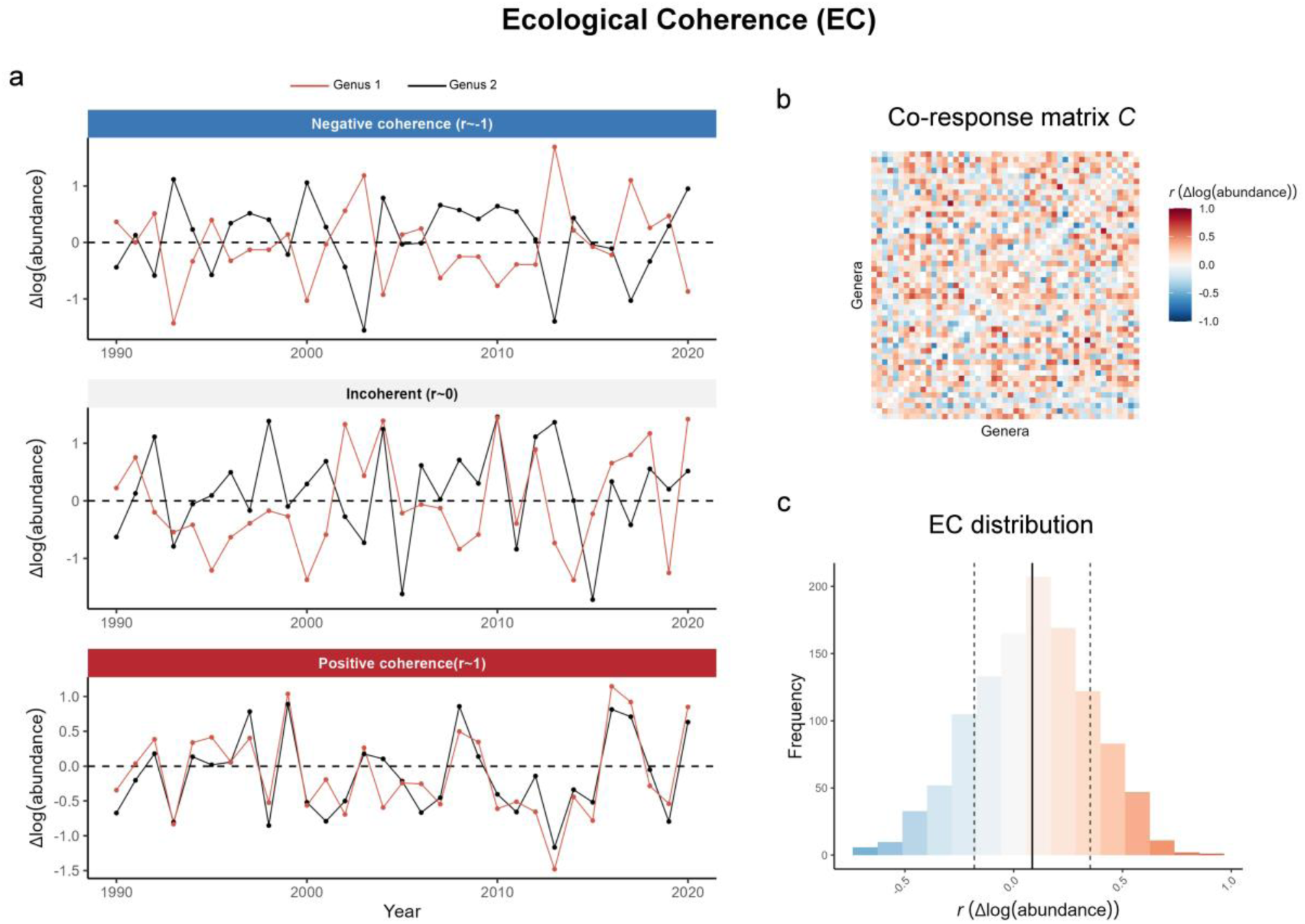
Conceptual illustration of Ecological Coherence (EC). EC captures the structure of co-responses within an assemblage and is expressed with two complementary statistical objects: a co-response matrix and the distribution of its values. For our study, we considered responses at the genus level, and defined them as year-to-year change in log abundance. (a) Two genera showing three coherence states: strongly negative (compensatory; *r* ∼ −1), incoherent (*r* ∼ 0), and strongly positive (*r* ∼ 1). Assemblages were defined as groups of genera co-occurring within spatial grid cells of 100 km for terrestrial regions and 500 km for marine regions. To compute EC within an assemblage, we considered the co-responses over a minimum of 10 common years between genus pairs. (b) The co-response matrix *C* is a square matrix where each value represents the pairwise correlation between taxa, providing insights into the structure of co-responses within an assemblage and the roles of individual taxa in shaping this structure. (c) The EC distribution is the distribution of correlations from the *C* matrix, and highlights the variability in direction and strength of coherence among taxa. The shape of this distribution serves as a profile that characterizes the EC of a given assemblage, facilitating its interpretation. The black vertical line indicates the mean, while the dashed vertical lines represent the standard deviation.

### Computing EC in empirical assemblages

#### Construction of assemblages

We used the BioTIME v1.0 dataset, the largest global dataset of raw abundance or biomass time-series data encompassing both marine and terrestrial systems (Dornelas et al. 2018). The dataset is organized by study and location (plot), and includes data spanning from 1874 to 2016. It was downloaded on 15 March 2023 (https://biotime.st-andrews.ac.uk/getFullDownload.php).

We grouped taxa co-occurring within spatial grid cells to delineate communities. There is no universally optimal grid size for representing the spatial distribution of diffuse taxa assemblages (Ricklefs 2008), and it depends on the study’s objectives. Excessively large scales may obscure ecologically meaningful interactions by conflating broad climatic responses with local contingencies. Conversely, overly small scales risk missing broader patterns of connectivity. To balance ecological relevance and spatial resolution, we chose a 100 km grid for terrestrial environments, consistent with prior global biodiversity studies (Blowes et al. 2019 using BioTime; Trisos et al. 2020), and a 500 km grid for marine environments, reflecting the larger ranges of marine taxa due to fewer geographical barriers, higher dispersal, and wide-ranging foraging behaviors (Rapoport 1994).

#### Yearly log-abundance change time series

We computed correlations between yearly changes in log-abundance for each taxon, following the methods in Lertzman-Lepofsky et al. (2024). For each assemblage, we first grouped the data to genus level, as species-level data did not reliably distinguish true zeros from low detection probabilities over time. We acknowledge that genus-level aggregation may both obscure genuine species-level relationships and introduce spurious ones. We then computed year-to-year log-transformed abundance changes for each species as Δ log(*abundance*) = log(*N*_*t*_) − log(*N*_*t*−1_), yielding a measure of proportional change that removes dependence on species’ abundance scale and facilitates comparability across taxa with different typical abundances and sampling scales (Collen et al. 2009, Leung et al. 2020, Loh et al. 2005). While differencing removes long-term trends and stabilizes variance, we note that biological autocorrelation (e.g., due to species life histories) and potential density dependence in population dynamics may still be present in the transformed series. Genera with fewer than 10 years of data were excluded, as previous research identified this threshold as the minimum for reliable trend detection (White 2019), and genera with zero variance in log-transformed abundance or biomass change were also removed. Finally, we did not consider temporal lags between taxa (i.e., delayed auto-correlations), as choosing the proper time scale would require additional assumptions and detailed data beyond the scope of this study.

#### Computation of EC

Once genera were grouped into assemblages and yearly changes in log abundance or biomass were calculated, we computed the co-response matrix by aggregating pairwise correlations between all genera. While numerous approaches exist for quantifying co-responses in time series, ranging from linear correlations to complex hierarchical models (e.g., Gouhier & Guichard 2014), we employed correlations to simplify analyses and ensure interpretability (standardized values from −1 to 1). Correlations for a given genus pair were calculated only for years common to both genera, with a minimum threshold of 10 overlapping years (Figure S2, S1, S5). This filtering ensured the reliability of the correlation estimates (Lertzman-Lepofsky et al. 2024). Values in the co-response matrix either represented valid correlation coefficients or were set to NA for genus pairs with fewer than 10 common years of data. Lastly, correlations exceeding 0.9 or falling below −0.9 were considered spurious and excluded, as extreme correlations in large, heterogeneous datasets may often reflect artefacts (e.g., shared absences, data anomalies) rather than true ecological signals, and it was not feasible to manually verify all pairs. Additionally, assemblages with fewer than 20 genera were discarded. Each assemblage’s EC was characterized by its co-response matrix (Figure S3) and the distribution of its elements (i.e., EC distribution) (Figure S4).

#### Regional or global summaries of EC distributions

We aimed to summarize EC distributions at regional and global scales. Because assemblages differ in species richness, their EC distributions cannot be directly compared. To make them comparable, we expressed each distribution as the percentage of pairwise correlations falling into predefined bins, rather than as raw counts. Specifically, correlation values were binned in increments of 0.1 (from −0.9 to 0.9) for each assemblage, and the percentage of pairwise correlations in each bin was calculated. We then visualized the distribution of these percentages across assemblages using boxplots, which capture both the central tendency and variability of correlation frequencies within bins. These summaries provide a broad-scale view of coherence structure at global (terrestrial vs. marine) and regional (ecoregion-level; see *EC distributions within ecoregions section*) scales.

### Null model approach to detect significant associations

An important challenge when assessing temporal coherence in community dynamics is to distinguish true ecological signals (e.g., shared responses to environmental drivers or taxa interactions) from spurious relationships that may occur at random. Correlations can be inflated or deflated in large datasets with relatively short or incomplete time series due to several factors, such as small sample size, shared missing observations, or low variance in one or both series. We used a null model approach based on randomization to generate a reference distribution for EC values expected under independence. To do so, we permuted the temporal order of measurements for each time series, thus preserving its marginal distribution (scale and variance) but removing any temporal and cross-taxa dependence. We repeated this procedure 1000 times, generating an ensemble of randomized co-response matrices *C* (EC values).

We compared the distribution of observed EC values to the distribution of randomized EC values using the Wasserstein distance, which provides a sensitive measure of dissimilarity between distributions (Figure S6). Specifically, we computed the Wasserstein distance of order p using the *wasserstein1d* function from the *transport* R package (Schuhmacher et al. 2024). This distance quantifies how different two distributions are, based on the cost of optimally “moving” one distribution to match the other. The parameter p controls how this cost is calculated: it sets how strongly large differences between values are weighted. When p = 1, all differences contribute linearly to the distance. When p = 2 or more, larger differences are emphasized more strongly. We tested both p = 1 and p = 2 and selected p = 2 for final analyses, as it provided greater sensitivity in distinguishing structured EC from the null distribution in our datasets, and placed greater emphasis on stronger correlations, which are ecologically more meaningful.

Finally, we computed the distribution of Wasserstein distances between observed vs. null EC and compared this to the distribution of null vs. null distances (computed by comparing pairs of randomized EC matrices). An empirical one-sided p-value was obtained for each assemblage as the fraction of null-vs-null distances exceeding the mean observed-vs-null distance. Assemblages were considered to exhibit non-random distributions of EC when p < 0.05.

### Variance partitioning of inter-genus coherence strength

We evaluated how EC is affected by taxonomic relatedness, hierarchical classification, and local community context. To do this, we modeled the variation in the absolute correlation of ecological responses between genus pairs within assemblages. We converted the genus-level correlation matrix for each assemblage to long format, where each row corresponds to a unique genus pair and its associated correlation value. We excluded self-pairs and missing values, and annotated each genus with its class identity. The resulting dataset linked each absolute correlation value to two genera (genus1, genus2), their classes (class1, class2), and the assemblage in which they co-occurred.

We partitioned the variation in EC using a linear mixed-effects model from the *lmer* function from the *lme4* R package (Bates et al. 2015), with EC as the response variable and crossed random intercepts for genus1, genus2, class1, class2, and assemblage. This structure partitions the total variance in pairwise ecological coherence into components attributable to genus-level identity, class-level affiliation, and assemblage context. The residual variance captures the remaining heterogeneity due to unmeasured variables.

### Genera contributions to EC

Within each assemblage, genera displayed varying degrees of correlation strength with others. We quantified contribution to Ecological Coherence with the absolute median of the absolute values in each column of the pairwise correlation matrix. This genus-level metric captures the typical strength of association a genus maintains with all other genera in the assemblage, regardless of the direction of the correlation. This allows us to identify genera that consistently exhibit strong correlations with others and may therefore play key roles in driving the coherence of the assemblage.. By summarizing genus-level contributions across assemblages, we characterized both the central tendency and variability of species’ roles in driving EC.

### EC distributions within ecoregions

We analyzed regional EC distribution summaries across different ecoregions, each characterized by distinct environmental conditions and taxonomic assemblages. Terrestrial assemblages were classified into biomes using the RESOLVE Ecoregions dataset (Dinerstein et al. 2017), while marine assemblages were assigned to realms based on the Marine Ecoregions of the World (MEOW) dataset from WWF (Spalding et al. 2007). Assemblages were linked to their respective biome or realm by intersecting their geographical coordinates with the spatial boundaries of these datasets. In cases where assemblages overlapped multiple biomes or realms (14 terrestrial and 32 marine intersected two, while one terrestrial and six marine intersected three), we applied a majority-rule assignment. We then summarized EC distributions at the biome and realm levels to capture the variability in co-responses across assemblages within each ecoregion.

### Sensitivity to sampling completeness and time series length

Sampling of an assemblage can directly influence EC and its interpretation. If sampling captures only a small, non-representative subset of an assemblage’s total taxonomic and functional diversity, the resulting EC may be biased. For instance, if only a specific lineage (e.g., a dominant clade of primary producers or closely related predators) is included, their shared ecological traits due to evolutionary relatedness may inflate the frequency of positive correlations in their responses (Johnson et al. 2024), leading to an artificially positively skewed EC distribution. At a broader spatial scale, comparing EC summaries of multiple assemblages within a terrestrial biome or marine realm introduces another layer of variability. Differences in the number of assemblages sampled per region can influence the EC, where regions with few assemblages may show higher variability compared to regions with larger sample sizes, which could tend to stabilize the EC variance.

Additionally, EC is sensitive to the length of the underlying time series, as has been shown for many variance-based metrics derived from population abundance data (Arkilanian et al. 2020). Shorter time series tend to produce less precise and more variable correlation values (Lertzman-Lepofsky et al. 2024), which can, in turn, influence EC. In fact, the BioTIME dataset is limited by short temporal spans across studies (Gonzalez et al. 2016). To mitigate this, we computed EC for pairs with at least 10 overlapping years of data. However, this threshold only partially addresses the issue: assemblages with generally shorter time series may still exhibit greater variability in correlation estimates, potentially distorting EC values.

We investigated the effect of sampling completeness with a series of additive linear models using the *lm* function in R. These models tested how taxonomic diversity, time-series length, and sampling effort influence the variance in EC among genera. The response variable was the variance of the EC distribution (referred to as EC variance). We first modeled EC variance at the assemblage level, fitting separate models for terrestrial and marine assemblages. The response variable was the EC variance within each assemblage. For each system, we fitted to alternative models: a model including genus richness and mean temporal overlap among genera (mean number of shared years), and a model including class richness and mean temporal overlap. We did not include genus and class richness in the same model because they were highly correlated. We also modeled EC variance at the ecoregion level, using the mean EC variance across assemblages within each terrestrial biome or marine realm as the response variable. Predictor variables included mean genus richness, mean class richness, mean temporal overlap among genera, and the number of assemblages per region. All variables were standardized using z-score transformation to allow direct comparison of effect sizes across predictors. We extracted standardized regression coefficients (β) along with their confidence intervals and significance levels, and assessed model fit using residual diagnostics.

## Results

### Consistent and contrasting EC distribution across assemblages

We assessed Ecological Coherence for 341 terrestrial and 105 marine assemblages. Assemblages whose EC distributions did not differ significantly from the null were excluded, resulting in 278 terrestrial and 104 marine assemblages retained for analysis (Figure 2ac). Terrestrial assemblages comprised, on average, 55.3 genera (SD = 23.7) and 1.5 classes (SD = 1.0), while marine assemblages included 78.7 genera (SD = 84.6) and 12.3 classes (SD = 10.1). The mean number of years shared between genera used to compute pairwise correlations was 21.2 years (SD = 2.9) for terrestrial assemblages and 14.4 years (SD = 5.5) for marine assemblages (Figure 2; Figure S1).

**Figure 2.**
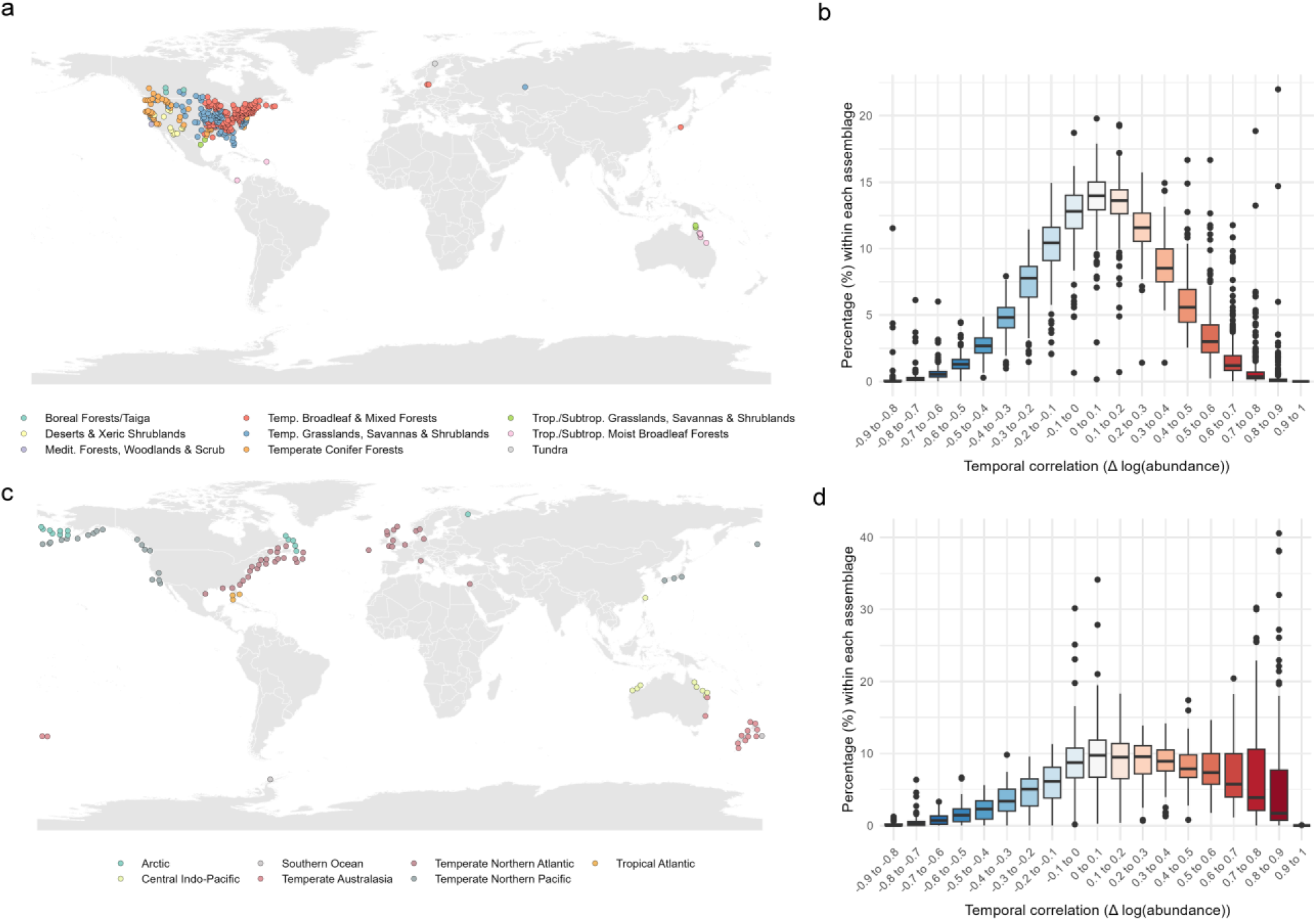
Ecological Coherence (EC) in natural assemblages. (a, c) Geographic distribution of the 278 terrestrial (a) and 104 marine (c) assemblages included in the analysis. These assemblages showed significant differences in EC relative to null model expectations. (b, d) Summary distributions of pairwise correlations across terrestrial (b) and marine (d) assemblages, representing the percentage of genus pairs falling within each EC correlation bin.

Despite substantial environmental and taxonomic differences, EC distributions showed a consistent structure with dominance of weak correlations. However, clear contrasts emerged between terrestrial and marine realms. Terrestrial assemblages, mostly composed of horizontally structured communities, exhibited near-normal EC distributions centered around zero, with few strong positive or negative correlations (Figure 2b). In contrast, marine assemblages—more taxonomically diverse and vertically structured—showed a pronounced skew toward positive correlations, including a higher frequency (a mean of 6.3%) of extreme positive correlations (between 0.8 and 0.9) (Figure 2d).

### Drivers of inter-genus coherence differ between terrestrial and marine systems

The crossed random effects model revealed contrasting drivers of variation in pairwise ecological coherence between terrestrial and marine assemblages. In terrestrial assemblages, genus identity accounted for a combined 20.2% of the variance, while class identity only explained a 1.9%. The assemblage factor accounted for 4.7% of the variance. In contrast, in marine assemblages, assemblage identity explained a much larger share of the variance (24.3%), while genus identity contributed 11.7%. Class-level contributions remained low (1.8% combined), similar to terrestrial systems. A substantial portion of the variance remained unexplained by the model (73.3% in terrestrial and 62.2% in marine assemblages), indicating heterogeneity in coherence not attributable to taxonomic identity or local context (Figure 3).

**Figure 3.**
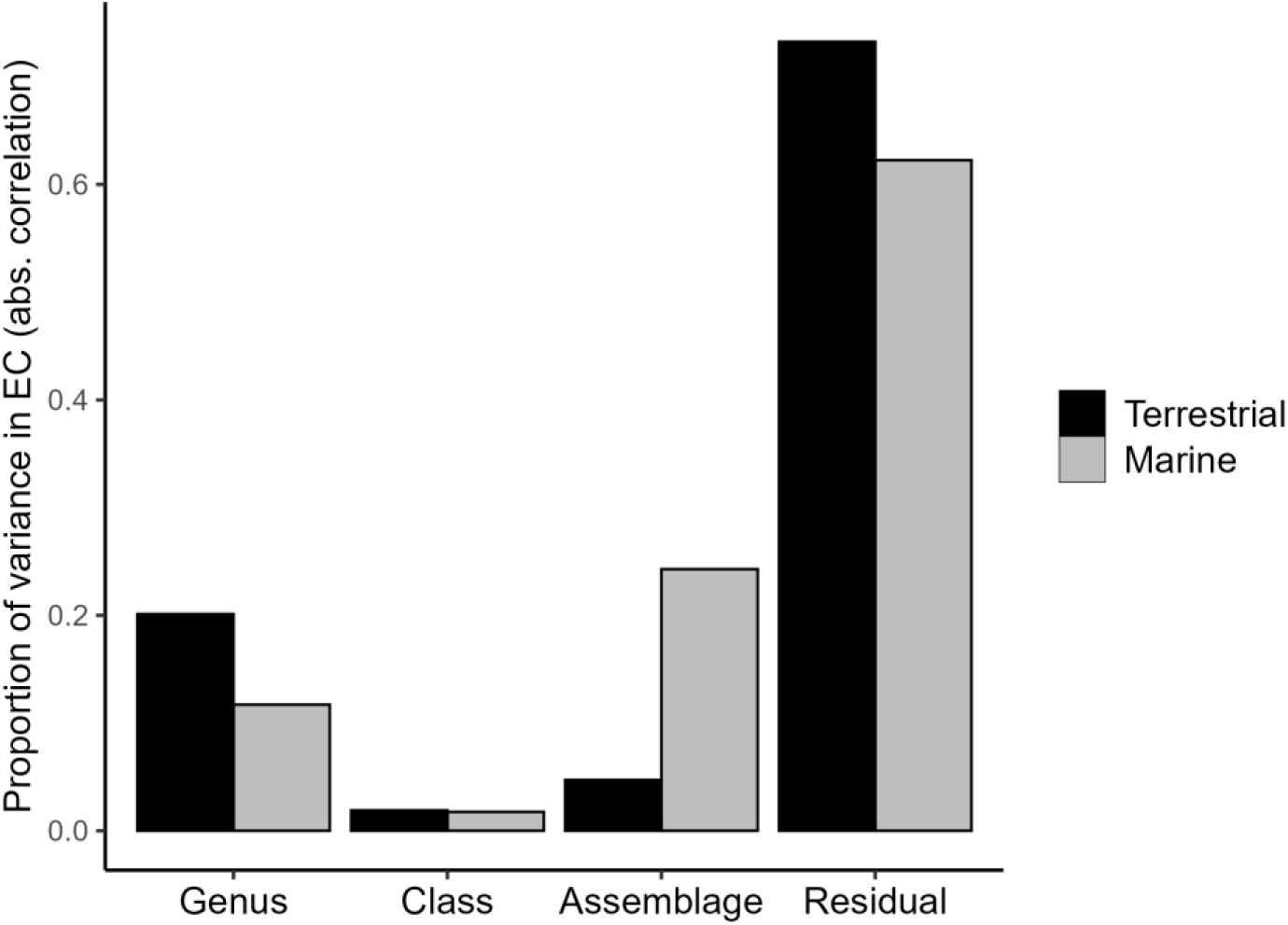
Proportion of variance in inter-genus ecological coherence explained by taxonomic and assemblage-level random effects. Variance components are based on a crossed random effects model fitted to the absolute correlation in ecological responses between genus pairs. Bars show the proportion of total variance attributable to genus identity, class identity, assemblage identity, and residual unexplained variation.

### A few genera dominate EC contributions across assemblages

Most genera showed small contribution values to overall EC, as measured by the median of the absolute values of their pairwise correlations with all other genera in the assemblage. Marine assemblages had higher mean genus contributions (0.35 versus 0.22) and variability (SD = 0.17 versus SD = 0.08). Outliers were widespread, with a small number of genera in many assemblages showing disproportionately strong contributions compared to others (Figure 4).

**Figure 4.**
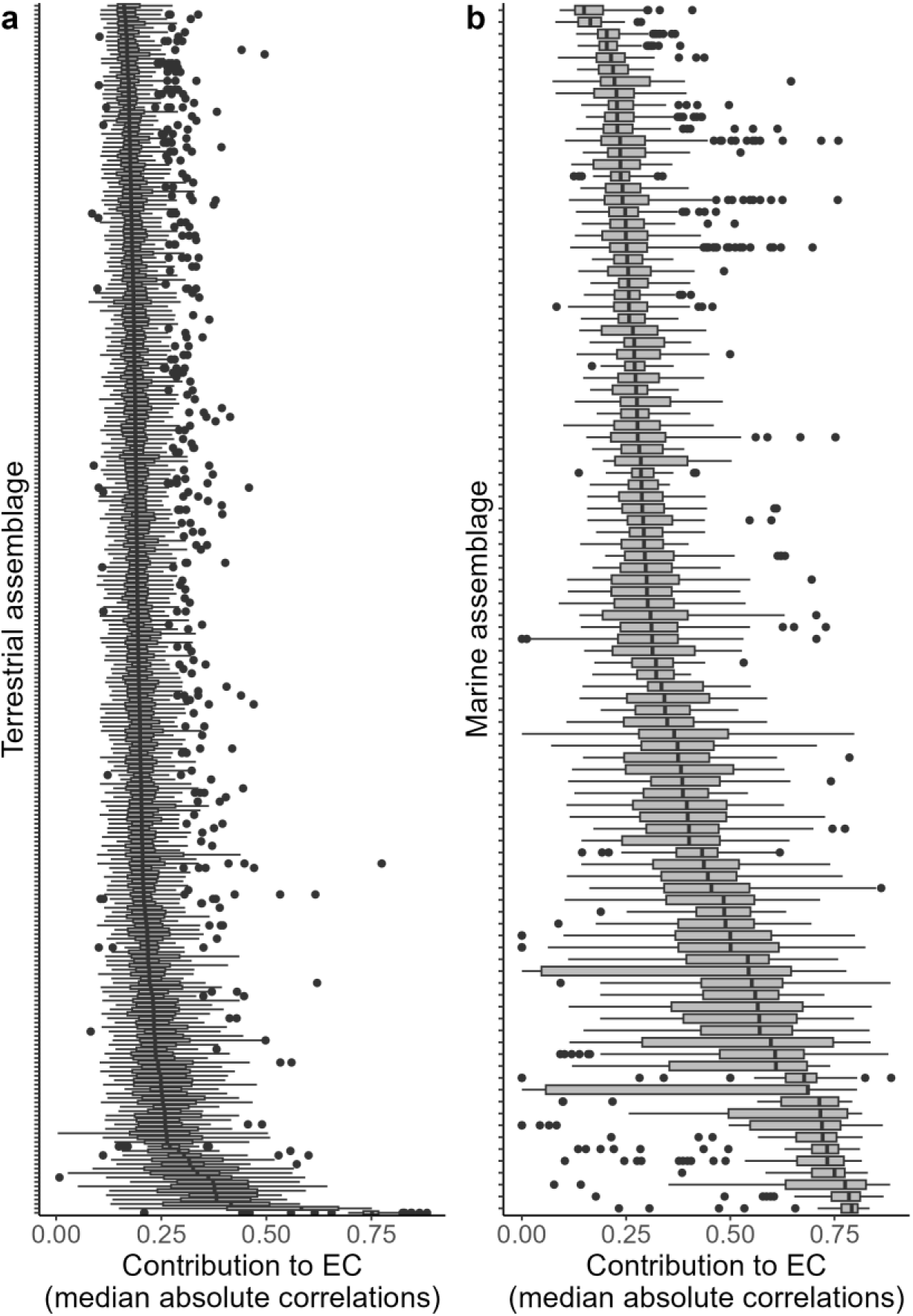
Genera contributions to Ecological Coherence (EC). The figure presents the distribution of genera contributions in (a) terrestrial and (b) marine assemblages. The contribution of a genus to EC within an assemblage is calculated as the absolute median of the absolute values of its pairwise correlations with all other genera in the assemblage. Higher values indicate that a genus typically exhibits stronger correlations—regardless of sign—with the rest of the community, suggesting a stronger role in shaping assemblage-level coherence.

### Sampling completeness

At the assemblage level, both genus-based and class-based models revealed consistent predictors of variance in EC. Longer temporal overlap between taxa was strongly associated with lower EC variance in terrestrial assemblages (β = –0.58, p < 0.001 in the model considering genus; β = –0.46, p < 0.001 in the model considering class). In contrast, taxonomic richness showed positive effects: both the number of genera (β = 0.20, p < 0.001) and the number of classes (β = 0.24, p < 0.001) significantly increased EC variance. These models explained ∼38% of the variance in EC (adjusted R² ≈ 0.38, F ≈ 82, p < 0.001).

In marine assemblages, the same negative effect of temporal overlap was detected (β = –0.33, p < 0.001 in both models). However, neither genus richness nor class richness had a significant effect (β = –0.12, p > 0.2 in both cases). The models explained a modest portion of the variance (adjusted R² ≈ 0.11), indicating lower explanatory power relative to terrestrial models.

Sampling completeness strongly influenced the variance in EC across terrestrial biomes. The model, which accounted for the effects of the number of assemblages summarized within each region, mean temporal depth of correlations, mean number of genera, and mean number of taxonomic classes per biome, explained 98.5% of the variance in mean EC variance (adjusted R² = 0.971, F = 67.5, p < 0.001). Among the predictors, the mean number of taxonomic classes had the strongest positive effect (β = 1.87, p < 0.001). The number of assemblages per biome (β = 0.683, p = 0.006) and the mean temporal depth of genera overlap (β = 0.591, p = 0.013) also had significant positive effects. Interestingly, the mean number of genera showed a significant negative effect (β = –0.321, p = 0.044) (Figure 5a).

**Figure 5.**
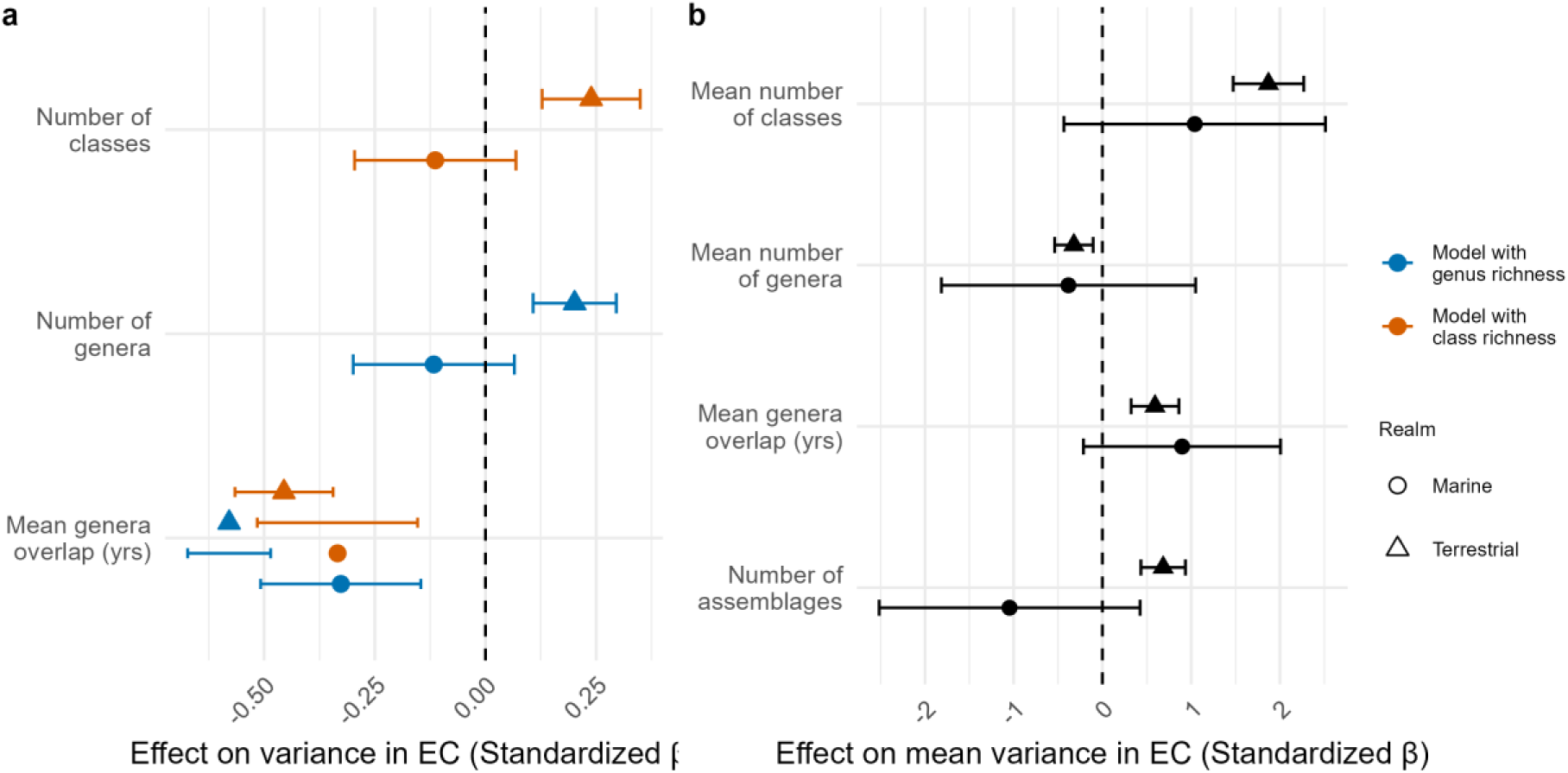
Effect of sampling completeness on Ecological Coherence (EC) at two spatial scales. (a) Standardized effect sizes (β) and 95% confidence intervals for predictors of within-assemblage EC variance, based on linear models fitted separately to terrestrial and marine assemblages. Each model includes either genus richness or class richness as a taxonomic predictor, along with the mean common years of overlap between genera. (b) Standardized effect sizes for predictors of mean EC variance across ecoregions (terrestrial biomes or marine realms), based on models including the number of assemblages, mean genera overlap, and mean genus and class richness.

In contrast, the model for marine realms explained 63% of the variance (adjusted R² = – 0.099, F = 0.856, p = 0.60), but none of the predictors were statistically significant.. Although effect sizes were comparable in magnitude to the terrestrial model, wide confidence intervals reflected greater uncertainty. Notably, the number of taxonomic classes had a positive effect (β = 1.04, p = 0.301), while the number of assemblages had a negative effect (β = –1.05, p = 0.297), mirroring the directions observed in the terrestrial model, though with insufficient statistical power to confirm these relationships (Figure 5b).

### Regional variations in EC

Specific ecoregions show strikingly different EC distributions. Tropical and subtropical moist broadleaf forests exhibited the highest frequencies of both strong positive (21.4%) and strong negative (10.6%) correlations, and a high proportion of extremely strong positive correlations (5.3% between 0.8 and 0.9). This contrasted with other terrestrial regions, where the proportion of strong correlations was generally below 10% and extreme correlations were rare (Figure 6a). The Central Indo-Pacific showed the most spread distribution, with over 60% of genus-level correlations exceeding 0.5, and more than 21.6% falling in the 0.8–0.9 range. Similarly, the Tropical Atlantic and Temperate Australasia also showed elevated levels of strong positive coherence (47.3% and 41.6%, respectively). In contrast, strong negative correlations were generally infrequent across marine regions, typically under 5% (Figure 6b).

**Figure 6.**
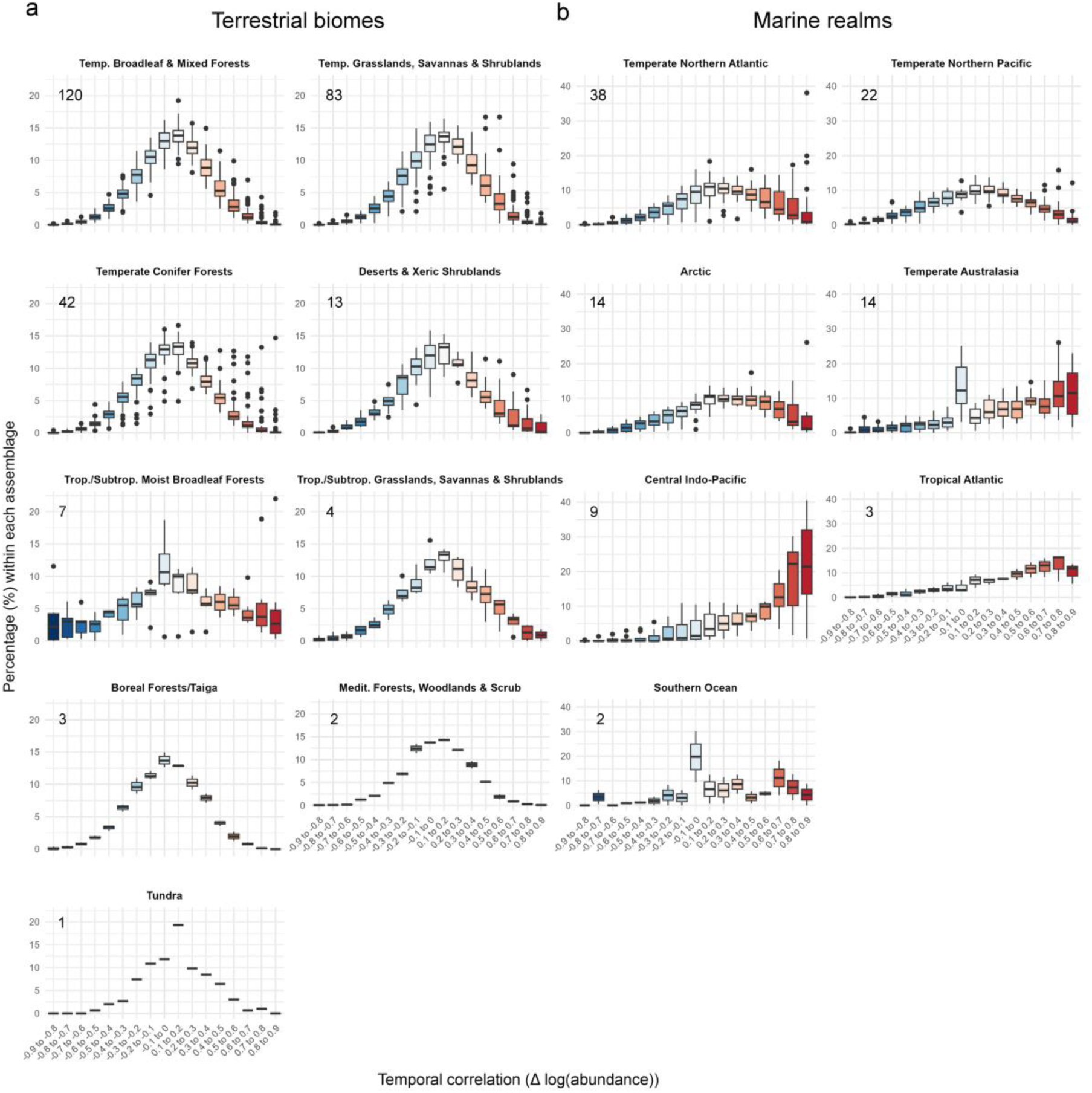
Summaries of assemblages’ Ecological Coherence distribution across ecoregions. The percentage of correlations observed within assemblages is shown by ecoregion for (a) terrestrial biomes and (b) marine realms. The number in the top left corner of each plot indicates the number of assemblages analyzed within each ecoregion.

## Discussion

Ecological Coherence (EC) provides a new framework for characterizing how co-responses between taxa are organized within communities. By combining the matrix of correlations among taxa with the distribution of these correlations, EC moves beyond single summary metrics of synchrony to capture both the structure and diversity of co-responses. Here, we computed EC from year-to-year changes in log-abundance, approximating per-capita growth rates and reflecting short-term fluctuations in population dynamics, across 446 assemblages worldwide. EC distributions revealed a general prevalence of weak correlations but also captured instances of strong, directional correlations. Moreover, analysis of the co-response matrix consistently identified a small subset of genera with strong correlations to many others, suggesting a promising path to detecting taxa that may play central roles in amplifying or buffering community responses to environmental change. Our findings raise questions about what drives coherence and how it shapes community response to environmental changes.

A first key finding is that most assemblages were dominated by weak correlations, which suggests that weak coherence may be an inherent feature of natural communities. Weak coherence may arise because taxa respond independently due to differences in their biologies, because they occupy heterogeneous microhabitats that expose them to distinct environmental drivers, or because ecological networks are sparse and many taxa do not interact strongly. Stronger coherence would therefore be expected only among subsets of taxa that are either linked by shared environmental filters or directly connected through biotic interactions (Lertzman-Lepofsky et al. 2025).

Beyond the dominance of weak coherence in EC distributions, we observed systematic differences between terrestrial and marine systems. Terrestrial assemblages, composed primarily of a few taxonomic classes within trophic guilds, showed normal-shaped EC distributions centered around zero. In contrast, marine assemblages, which are more taxonomically diverse and trophically structured, exhibited a much more skewed distribution, towards strong positive correlations. This contrast may reflect complementary explanations.

First, in more vertically structured communities, energy flows across trophic levels may amplify dependencies among species, thereby promoting stronger coherence in their responses. In contrast, terrestrial assemblages, mostly composed of horizontally structured communities dominated by primary producers, may experience more competition, leading to greater prevalence of negative or compensatory dynamics. Second, terrestrial assemblages often encompass fine-scale habitat heterogeneity—from dry, mesic, and humid habitats to grasslands and forests within a few hundred meters—which can decouple species responses. By contrast, marine assemblages are embedded in more homogeneous environments, where species experience similar conditions. Third, marine communities may also be more strongly and uniformly affected by large-scale environmental perturbations and physical connectivity. Broad-scale drivers, coupled with larval dispersal and the mobility of wide-ranging species, can synchronize responses across taxa and regions (Liebhold et al. 2004, Houlahan et al. 2007, Gouhier et al. 2010, Valencia et al. 2022). For example, Granzotti et al. (2024) showed that environmental drivers in a multi-trophic fish community explained much of the observed synchrony. These observations raise a broader question: what are the consequences of these different distributional shapes for community stability?

Previous studies have linked strong positive coherence to instability under anthropogenic pressures, with extreme events, overfishing, and environmental change inducing strong synchrony that can precede ecosystem collapse (Keitt 2008, Koenig & Liebhold 2016, Pedersen et al. 2017, Arimitsu et al. 2021). Conversely, negative coherence, often associated with compensatory dynamics, can stabilize communities by dampening aggregate variability (Gonzalez & Loreau 2009), though this effect may be limited to functionally similar or monotrophic systems. In multi-trophic communities, strong negative coherence may instead reflect divergent responses across functional groups, potentially contributing to instability (Thackeray et al. 2010, Viviani et al. 2019). In complex communities with multiple trophic levels, it is probable that both ends of the coherence spectrum—strongly positive and strongly negative—may signal instability, depending on trophic structure and functional diversity (Huang et al. 2020, Danet et al. 2021). From this perspective, the dominance of weak coherence in our global assemblage data may point to a stabilizing structure: weak interspecific covariance could buffer cascading effects and mitigate community-level responses to perturbations. This hypothesis highlights the importance of not only the direction but the magnitude of coherence in shaping ecological resilience. Furthermore, because marine assemblages encompass more trophic levels than terrestrial ones, their tendency toward more positive coherence may reflect the maintenance of consumer-resource coupling and network architecture (Boyce et al. 2015). Future work under the EC framework should investigate whether EC distributions skewed toward extreme correlations signal greater ecological vulnerability, either due to current environmental pressures or as lasting signatures of past disturbances.

We next examined the sources of variation in coherence across taxa.n terrestrial ecosystems, taxonomic identity (particularly at the genus level) is a stronger determinant of inter-genus coherence, while in marine ecosystems, local community context plays a more dominant role. In both systems, class-level identity contributes only marginally to explaining coherence variation, reinforcing the idea that coherence emerges from finer-scale taxonomic or ecological relationships.

Yet a substantial portion of the variance in coherence remained unexplained in both systems. This unexplained variance may reflect species-specific responses within genera, stochastic ecological processes such as disturbance events, or unmeasured environmental factors such as microclimate variation, habitat structure, or higher-order effects of the network of biotic interactions. Additionally, methodological noise or mismatches in temporal resolution could also contribute.

We further identified key taxa that disproportionately shape EC. Our analysis of genera contributions to EC revealed that, in nearly all assemblages, a few taxa exhibited disproportionately high contributions compared to the rest. This suggests that environmental perturbations affecting these key taxa could trigger strong cascading responses across the community, particularly for taxa that covary with them. Such observations may be expected for ecosystem engineers—such as corals, mangroves, or seagrasses—whose structural roles support numerous other taxa. Similarly, dominant taxa that drive ecosystem functioning, such as keystone tree taxa, seed dispersers, or top predators, may also exhibit high EC contributions due to their central ecological roles. If a taxon with a strong contribution to EC also occupies a central position in the network structure, it could have a heightened potential to drive widespread shifts in community dynamics and ecosystem functioning. Pedersen et al. (2017) documented strong covariance in abundance dynamics in response to overfishing before the collapse of a marine food web, with cod—the most central species in the network—exhibiting the highest contribution to coherence during this period (Fuster-Calvo et al. *in prep*). This highlights the potential for EC contributions to serve as an early indicator of taxa that play outsized roles in stabilizing or destabilizing communities. Further research should explore the EC contributions of different taxa and their structural positions in ecological networks, as this could provide valuable insights into identifying key taxa that mediate community-wide responses to environmental change.

Regional analyses revealed additional contrasts in EC distributions. Among terrestrial ecoregions, tropical and subtropical forests stood out for their broader EC distributions, characterized by a higher frequency of both extreme positive and negative correlations than any other terrestrial region. Similarly, in marine systems, the Central Indo-Pacific shows the highest proportion of extreme positive correlations of all ecoregions analyzed. A common feature of these regions is their exceptionally high taxonomic and functional diversity, spanning a wide range of trophic roles. Tropical forest assemblages, for example, include diverse plant groups (angiosperms, gymnosperms, ferns), arthropods (insects, spiders, crustaceans), and vertebrates (birds), while Indo-Pacific marine assemblages feature reef-associated and pelagic fish, elasmobranchs (sharks and rays), and various crustaceans. Such high diversity in trophic groups may broaden the range of responses within communities, amplifying both positive and negative correlations and leading to wider EC distributions, consistent with our earlier hypothesis for explaining terrestrial–marine differences in EC.

Another plausible explanation is that these distributions are further shaped by anthropogenic pressures and climate-driven disturbances. Both tropical forests and Indo-Pacific marine systems are subject to intense anthropogenic pressures and are especially vulnerable to climate-driven disturbances (e.g., warming, habitat degradation, coral bleaching), which may enhance synchrony or divergence in species responses, thereby amplifying EC variance (Kharouba et al. 2018, Song et al. 2024).

Finally, we investigated how taxonomic structure and temporal depth influence EC variance across scales. Taxonomic diversity consistently influenced variation in pairwise correlations among genera (EC variance). At the assemblage level, greater genus or class richness was associated with higher EC variance in terrestrial systems, suggesting it broadens taxa’ responses and types of interactions. In contrast, marine assemblages—already richer in taxa—showed no effect of richness on EC variance, which may reflect a saturation, where added taxa contribute little new functional variation (Valencia et al. 2022).

At the ecoregion level, class richness again increased EC variance in terrestrial biomes, while genus richness showed the opposite effect.. This discrepancy may reflect that adding more genera, while holding higher-level diversity constant, introduces more similar taxa, diluting variation. Alternatively, regions with many genera within the same classes or trophic levels may show reduced contrasts in responses, since added genera contribute redundancy rather than new functional roles..

Temporal structure also showed scale-dependent effects. At the assemblage level, longer temporal overlap among genera was consistently associated with lower EC variance in both realms, likely because longer time series may better capture stable co-responses and smooth short-term fluctuations (Arklianian et al. 2020). At the ecoregion level, however, greater temporal depth increased EC variance, perhaps because assemblages span different environmental conditions, histories, or disturbance regimes.

The number of assemblages per region also mattered.. In terrestrial biomes, regions with more assemblages exhibited greater EC variance, likely because they span broader environmental gradients and encompass more diverse community types. In marine realms, the effect was negative but non-significant, possibly reflecting the limited sample size or the greater environmental homogeneity of marine systems.

Finally, an additional sampling bias comes from the fact that our analysis is inherently constrained by the spatial and temporal biases of global ecological datasets (Gonzalez et al. 2016). The data are concentrated in well-studied locations, particularly in regions with extensive long-term monitoring or direct human impacts, while many ecosystems—such as remote terrestrial biomes, tropical areas and large portions of the Global South, and relatively undisturbed areas—remain underrepresented. Furthermore, longer time series are needed to detect ecological trends that unfold gradually or exhibit delayed responses to environmental change.

### Future work on the drivers and consequences of EC

#### Drivers of EC

The correlated variation between species’ abundances and occurrences across environmental gradients contains a wealth of information about the abiotic and biotic drivers that drive multiple species’ abundances and distributions to vary through space and or through time. To investigate the drivers of EC, it is essential to first model species’ responses in the context of their assemblage through hierarchical modelling approaches, and most importantly, to explicitly capture this correlated variation within this multispecies model. To achieve this, we can build on advances in joint species distribution models (Pollock et al. 2014) and hierarchical modeling of species communities (Ovaskainen & Soininen 2011) to build hierarchical models that explicitly estimate the covariance between species. One such way of doing so is to use latent variable models to capture species’ temporal or spatial correlations to latent variables (Warton et al. 2015), which capture hidden signatures of unmeasured predictors such as species interactions or unmeasured environmental gradients (Ovaskainen et al. 2016). This latent variable modelling approach yields a covariance matrix describing species’ co-responses (Warton et al. 2015), which can be used to investigate EC, and an estimation of common latent trends across species, which can be leveraged to explore the temporal or spatial gradients driving multi-species dynamics (Rigal & Knape 2023). To investigate the drivers of the species’ trends and the community EC, these estimated latent trends can then be compared to potential drivers of change to look for potential spatial or temporal matches with potential drivers of change. Specifically, similarities in the timing or spatial variation between the latent trends and potential drivers of change could hint at the possibility of attributing community-wide dynamics to these drivers with deeper exploration and appropriate data. Groups of species with similar dynamics can also be identified to understand how these drivers influence some species more strongly than others, and potentially which species’ responses scale up most strongly to influence community dynamics (Rigal & Knape 2023).

#### EC and ecological networks

Interactions ultimately shape how EC influences community dynamics, either amplifying or dampening its effects (Ives et al. 2000, 2003). While EC describes the distribution of species co-responses within a community, its ecological consequences will be better understood when examined in the context of coherence within and among trophic guilds and functional modules (Viviani et al. 2019). For instance, some studies suggest that synchrony among primary producers may have a stronger impact on entire food webs than synchrony within other trophic groups (Arimitsu et al. 2021, Zhao et al. 2023), and Rao et al. (2024), using phytoplankton-phytoplankton communities, show that community asynchrony within guilds increases stability, but between guilds could destabilise consumer communities.

With the increasing availability of large-scale ecological interaction databases (e.g., Albouy et al. 2019, Maiorano et al. 2020) and advancements in interaction inference methods (Poisot et al. 2015, Strydom et al. 2021), it is becoming more feasible to approximate network structures across multiple assemblages. The coherence in multi-trophic communities is only recently being investigated (Vinebrooke et al. 2004, Long et al. 2011, Albouy et al. 2019, Maiorano et al. 2020, Firkowski et al. 2022, Siqueira et al. 2024). The EC framework introduced here provides an opportunity to integrate interaction data, allowing us to examine coherence not just across entire communities, but within and among different network structures, such as trophic guilds or functional modules. This approach also strengthens the link between empirical and theoretical research, as models investigating the role of EC in community dynamics and ecosystem functioning can be grounded in concrete empirical structures—specifically, the co-response matrix and its distribution. Ultimately, integrating EC with network structure could pave the way for using these empirical distributions as indicators to better understand the mechanisms driving community responses to environmental changes and to anticipate potential ecological disruptions.

## Acknowledgements

Financial support was provided by the NSERC - CREATE Training program in computational biodiversity science and the NSERC Discovery Grant to DG.

