## Supplement 1 for "Ecological coherence in abundance dynamics across terrestrial and marine assemblages"

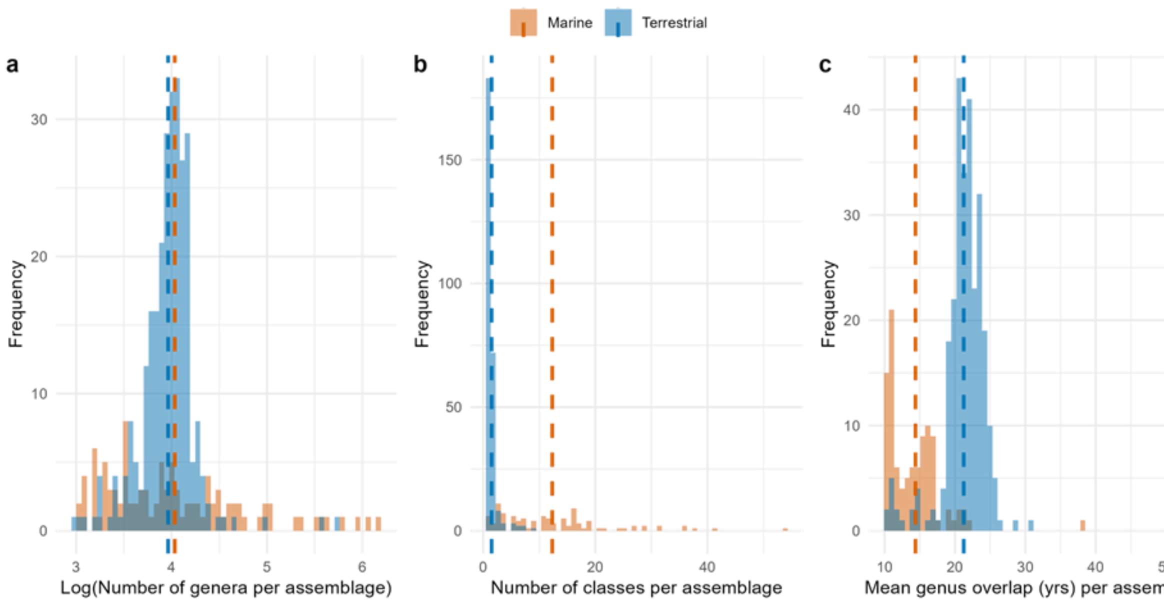


**Figure S1. Data distribution of terrestrial and marine assemblages.**

Dashed lines indicate the mean.

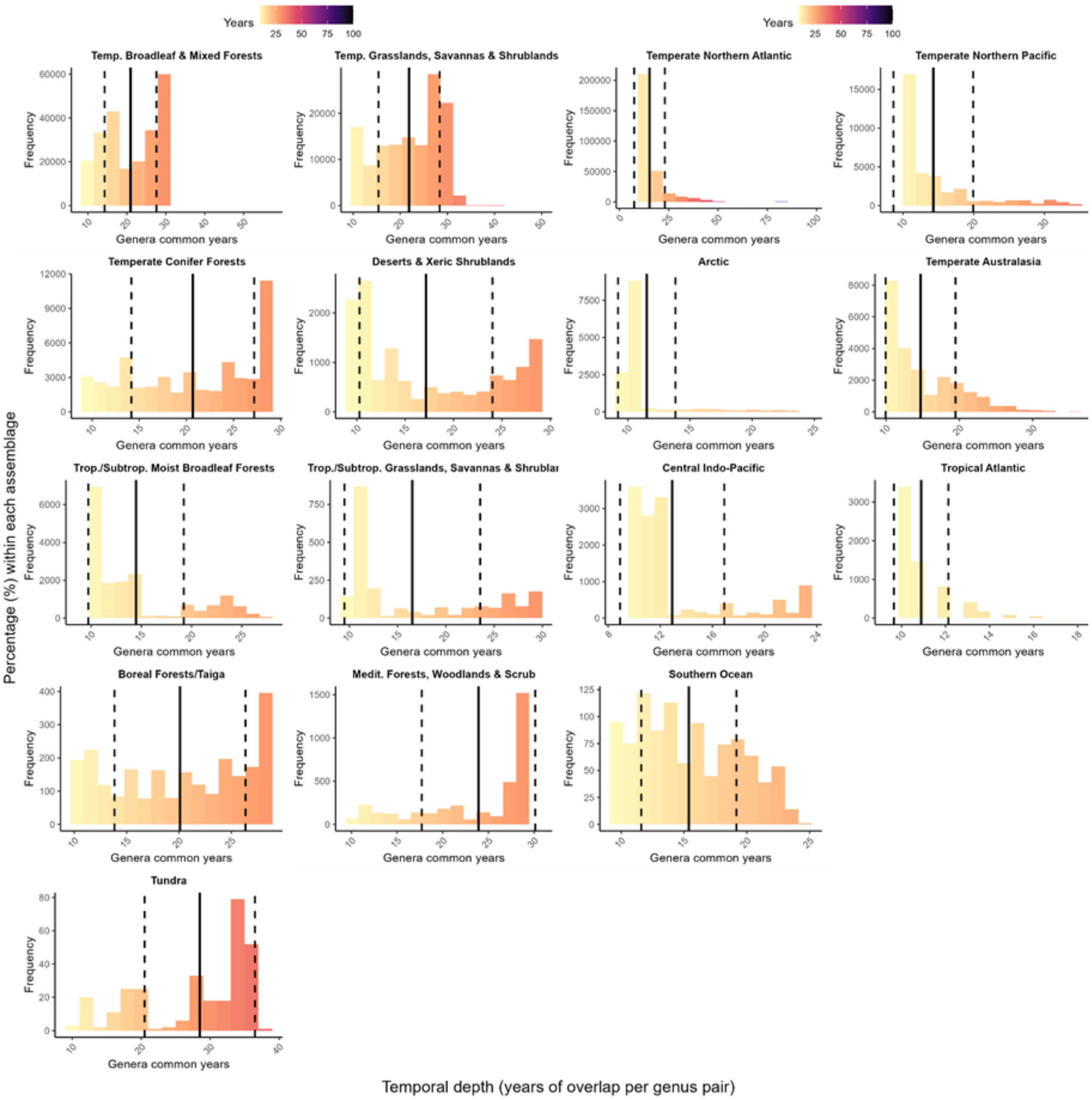


**Figure S2. Genera common years used to compute correlations in assemblages within each terrestrial biome and marine realm.**

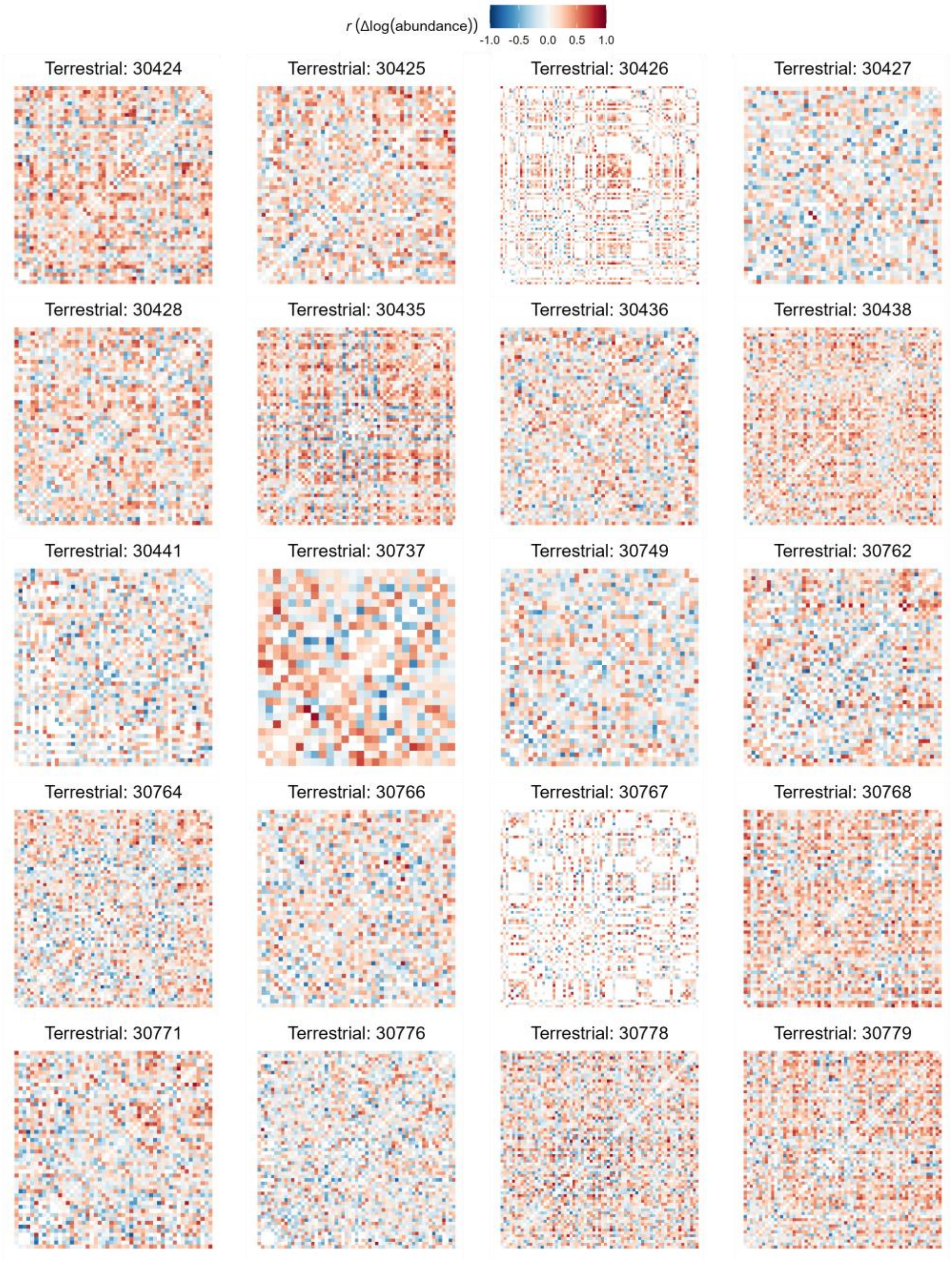


**Figure S3. EC co-response matrices for a set of terrestrial assemblages.**

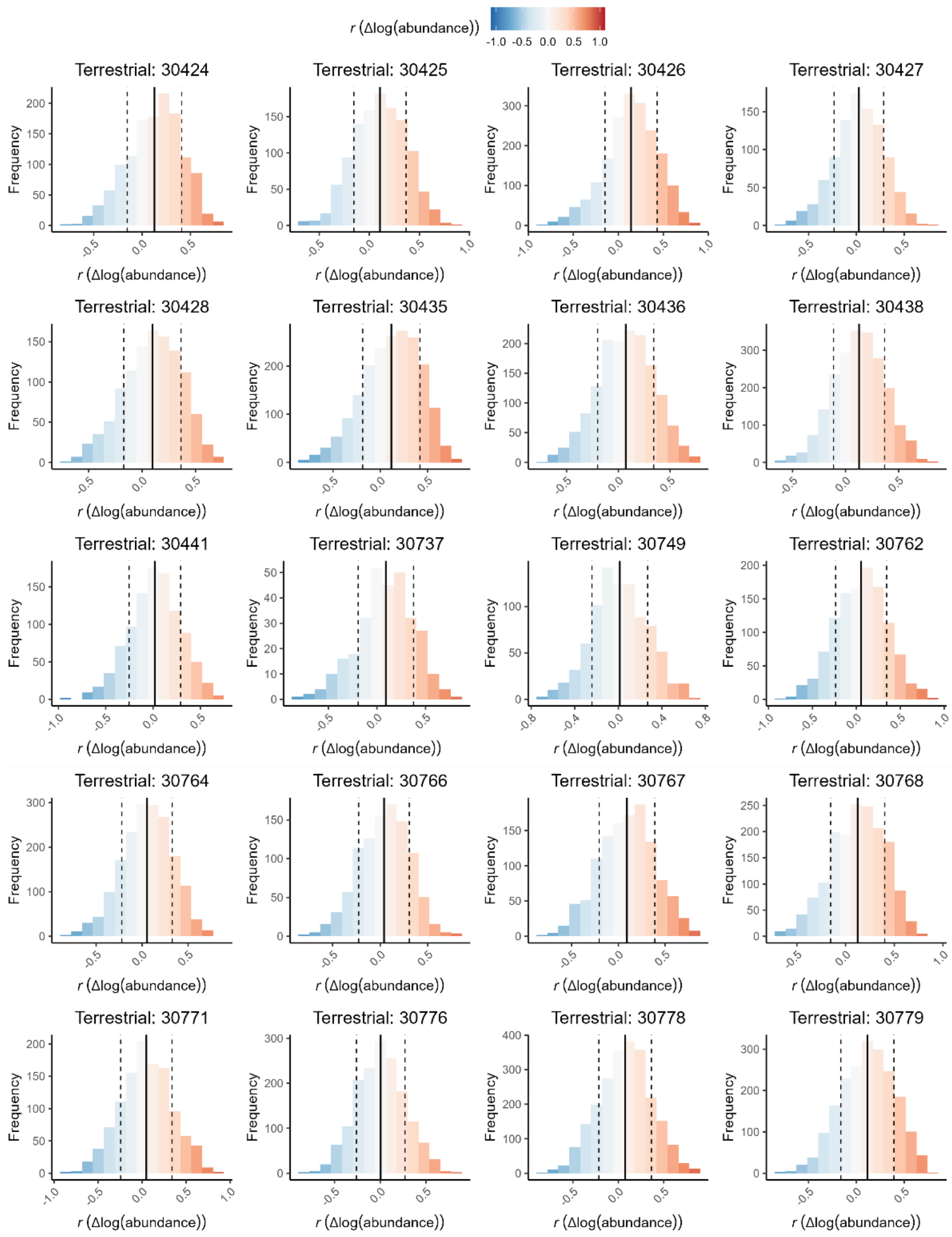


**Figure S4. EC distributions for a set of terrestrial assemblages.**

The black vertical line indicates the mean, while the dashed vertical lines represent the standard deviation.

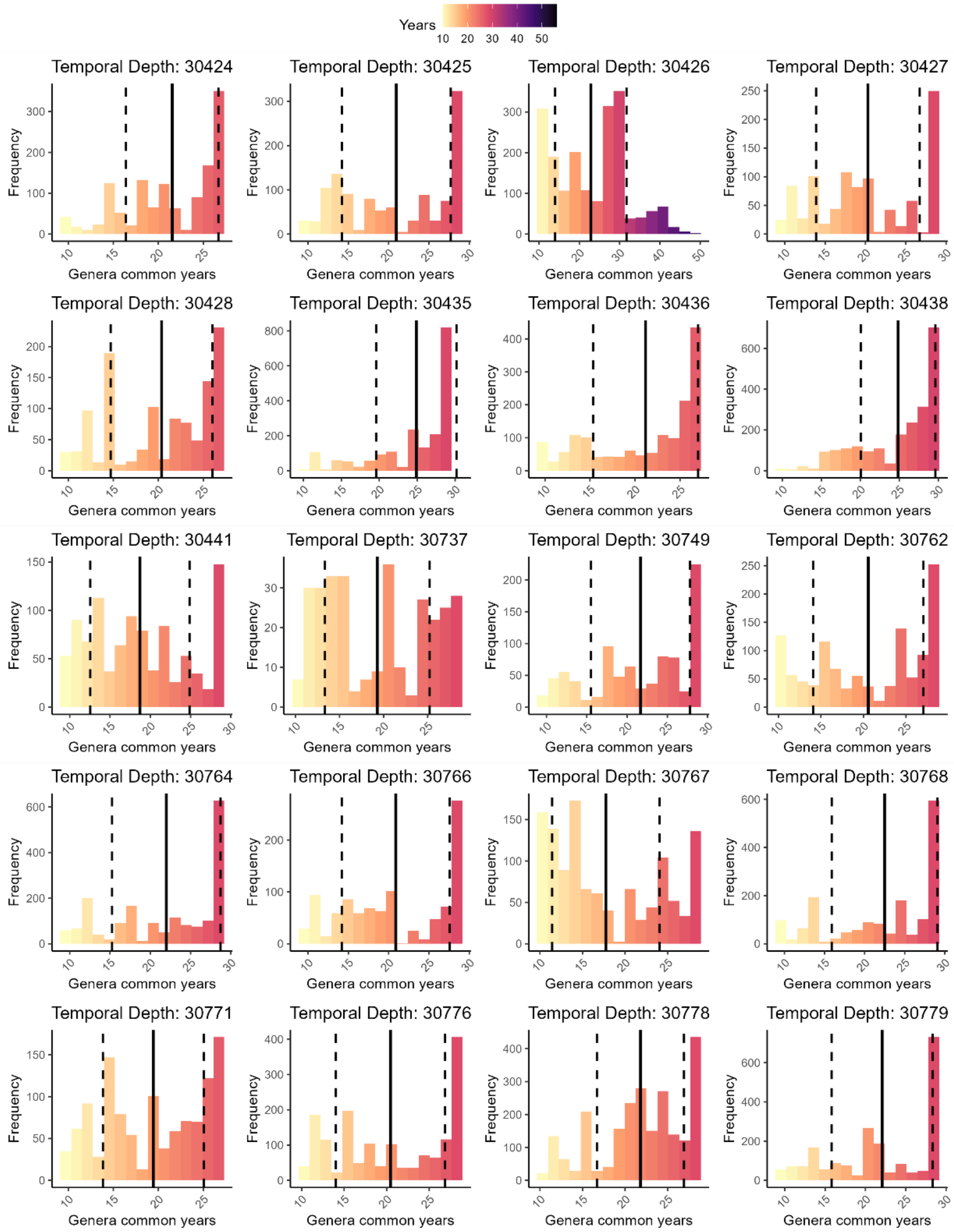


**Figure S5. Temporal depth (number of common years) of pairwise time series correlations for a set of terrestrial assemblages.**

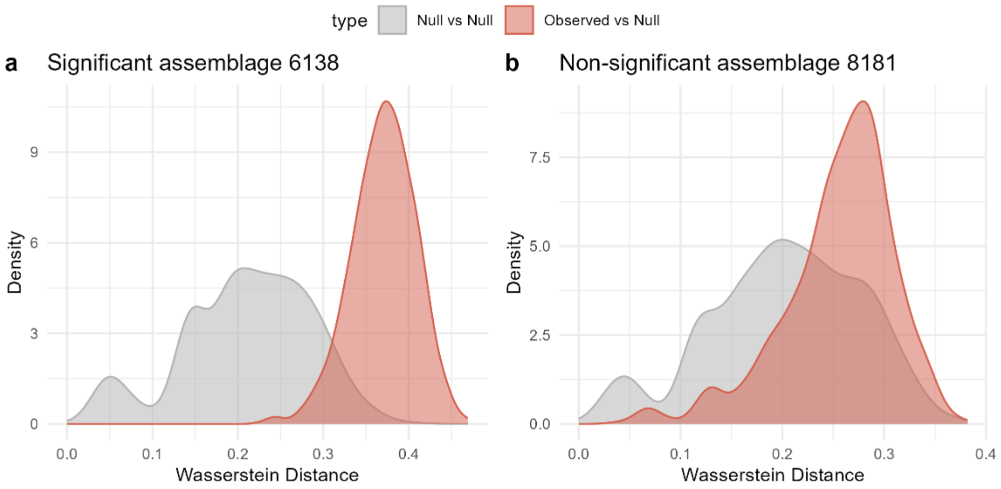


**Figure S6. Example assemblages illustrating tests of Ecological Coherence (EC) distributions against null expectations.**

For each assemblage, we computed a distribution of distances between observed and null EC (red) and a reference distribution of distances between pairs of null EC matrices (grey). Significance was assessed by comparing the mean of the observed–null distribution against the null–null distribution. The left panel shows an assemblage with significantly different distributions ($p<0.05$), indicating non-random EC structure, while the right panel shows a non-significant case consistent with random expectations.
